# Introgression from *Oryza meridionalis* into domesticated rice *Oryza sativa* results in shoot-based iron tolerance

**DOI:** 10.1101/2020.06.05.135947

**Authors:** Andriele Wairich, Ben Hur Neves de Oliveira, Lin-Bo Wu, Varunseelan Murugaiyan, Marcia Margis-Pinheiro, Janette Palma Fett, Felipe Klein Ricachenevsky, Michael Frei

## Abstract

Iron (Fe) toxicity is one of the most common mineral disorders affecting rice (*Oryza sativa*) production in flooded lowland fields. *Oryza meridionalis* is endemic from Northern Australia and grows in regions with Fe rich soils, making it a candidate for use in adaptive breeding. Aiming to understand tolerance mechanisms in rice, we screened a population of interspecific introgression lines (IL) from a cross between *O. sativa* and *O. meridionalis* for the identification of QTLs contributing to Fe toxicity tolerance. Six putative QTLs were identified. A line carrying one introgression from *O. meridionalis* on chromosome 9 associated with one QTL was highly tolerant despite very high shoot Fe concentrations. Physiological, biochemical, ionomic and transcriptomic analyses showed that the IL tolerance could partly be explained by Fe retention in the leaf sheath and culm. We constructed the interspecific hybrid genome *in silico* for transcriptomic analysis, and identified differentially regulated introgressed genes from *O. meridionalis* that could be involved in shoot-based Fe tolerance, such as metallothioneins, glutathione S-transferases and transporters from ABC and MFS families. This is the first work to demonstrate that introgressions of *O. meridionalis* into the *O. sativa* genome can confer increased tolerance to excess Fe.

**Highlight:** We identified QTLs associated with iron tolerance derived from *O. meridionalis*, and characterized their physiological basis in *O. sativa*.

## Introduction

Iron (Fe) is an essential microelement for almost all living organisms. In plants, Fe plays a critical role in respiration and photosynthesis and functions as a co-factor of many enzymes. However, when taken up excessively by plants, Fe becomes toxic as it participates in Fenton reactions catalyzing the generation of reactive oxygen species (ROS) (Guerinot and Yi, 1994). Fe toxicity (+Fe) is one of the most common mineral disorders affecting rice (*Oryza sativa* L.) production and yield in flooded lowland fields, where reducing conditions lead to excess availability of soluble ferrous iron (Becker and Asch, 2005, Fageria and Rabelo, 1987).

Fe toxicity symptoms include leaf bronzing, reduced tillering and poor development of roots, which become coated with a dark brown layer of Fe named Fe plaque (Becker and Asch, 2005, Audebert, 2006, Zhang *et al*., 2016). However, different shoot or root-based mechanisms can confer tolerance to +Fe (Becker and Asch, 2005): (1) formation of root plaque through the precipitation of oxidized Fe(III), which acts as a physical barrier and prevents excessive Fe uptake; (2) retaining Fe in ‘dumping sites’, at which Fe is immobilized and deposited in a metabolically inactive form in root tissue (Becker and Asch, 2005, Audebert and Sahrawat, 2000); (3) shoot-based tolerance mechanism associated with the retention of Fe in less photosynthetically active tissues such as leaf sheath or culm, reducing the impact on photosynthesis; or on the storage of Fe excess in the apoplasm, vacuole and in plastids (by ferritin adsorption) (Engel *et al*., 2012, Wu *et al*., 2019, Silveira *et al*., 2009); (4) control of Fenton-induced ROS formation by modulating plant redox homeostasis (Wu *et al*., 2017).

Yield losses caused by +Fe usually range between 15% and 30%, but can reach the level of complete crop failure, depending on the intensity of the stress and plant age (Becker and Asch, 2005, Asch *et al*., 2005). Strategies for mitigating yield losses include the selection of tolerant genotypes (Audebert, 2006). However, Fe tolerance is a quantitative trait controlled by multiple *loci* and genes. Moreover, the response of each specific genotype is dependent on its interaction with the environment (Dufey *et al*., 2010, Wu *et al*., 2014).

A significant constraint to the development of modern and high-yielding varieties is that the genetic diversity of the species *O. sativa* has been narrowed during rice domestication, and many traits associated with tolerance to adverse conditions (biotic or abiotic) may have been lost (Zhu *et al*., 2007, Palmgren *et al*., 2015, Menguer *et al*., 2017). On the other hand, abundant natural variation in landraces and crop wild relatives has been conserved in gene banks and constitutes a valuable genetic reservoir that can be used for crop improvement. Thus rice wild relatives (RWR) have particular importance as donors of genetic variation for cultivated rice (Arbelaez *et al*., 2015, Menguer *et al*., 2017, Stein *et al*., 2018).

Among RWR species, *O. meridionalis* is a promising source of tolerance to +Fe. A screening of several genotypes representing 21 species of the genus *Oryza* for Fe stress tolerance found two tolerant accessions of *O. meridionalis* (Bierschenk *et al*., 2020). Tolerance to high Fe concentrations is likely related to the origin of the species, which evolved in Northern Australia, at the edges of freshwater lagoons, seasonal wetlands and swamps, in addition to Fe rich soils (Vaughan, 1994, Juliano *et al*., 2005). *O. meridionalis* is part of the primary gene pool (wild species closely related to *O. sativa*), all containing AA genomes, which can cross breed for adaptive rice breeding (Sotowa *et al*., 2013, Bierschenk *et al*., 2020).

This study aimed at deciphering interspecific genetic variation in adaptation to +Fe and the underlying physiological mechanisms. This is a timely approach because powerful resources for the exploitation of the global genetic diversity of the genus *Oryza* have been developed recently, including populations derived from interspecific crosses (Arbelaez *et al*., 2015), as well as genome sequences of a variety of RWR including *O. meridionalis* (Stein *et al*., 2018). We used previously developed introgression lines (IL) generated by crossing *O. sativa* and *O. meridionalis* and screened it for tolerance to +Fe. Our specific hypotheses were (i) novel tolerance QTL for +Fe can be identified by crossing the species barrier of *O. sativa;* (ii) *O. meridionalis* introgressions can contribute novel adaptive traits that have not previously been described in *O. sativa;* and (iii) transcriptome analyses using an *in silico* constructed hybrid genome can shed light on the mechanisms underlying tolerance in an interspecific hybrid.

## Materials and Methods

### Plant material and screening for +Fe tolerance

A population of 32 interspecific ILs was used in a screening for tolerance to +Fe, which was generated from crosses between *O. sativa* cv Curinga (hereafter “*O. sativa*”) and *O. meridionalis* Ng. accession W2112 (Arbelaez *et al*., 2015). Seeds were originally obtained from Prof. Susan McCouch (Department of Plant Breeding and Genetics, Cornell University).

Experiments were conducted in a climate-controlled greenhouse in Bonn, Germany, from July to September 2018 (1^st^ experiment) and from March to May 2019 (2^nd^ experiment). Natural light was supplemented with artificial light between 7 am and 9 pm to ensure a minimum photosynthetic photon flux density of 400 μmol m^−2^ s^−1^. The minimum day/night temperature was set to 28/22 °C. Seeds were exposed to 55 °C for three days to break dormancy. Seeds were germinated and seedlings were grown in a mesh floating on solutions containing 0.5 mM CaCl_2_ and 10 µM FeCl_3_ for two weeks at 28 °C. The solutions were replaced twice a week. Eight seedlings per genotype per treatment (n = 4) were transplanted into 60 L containers (totalizing 8 containers), each containing 40 seedlings, filled with quarter-strength nutrient solution (Yoshida *et al*., 1976). After one week, nutrient solutions were changed to half-strength, and thereafter replaced with a full-strength solution once a week (Wu *et al*., 2019). The pH was adjusted to 5.5 every two days.

After three weeks of growth, plants were exposed to 250 mg L^-1^ of Fe as FeSO_4_.7H_2_O for fifteen days. Nitrogen gas was percolated into the culture solutions for 15 minutes every 4 hours. The reflectance between 380 - 1050 nm of the three last fully expanded leaves was determined after 1, 3, 5, 7 and 13 days after the onset of treatment, with PolyPen RP 410 (Photon Systems Instruments Drasov, Czech Republic). Leaf bronzing scores (LBS) were attributed to the three youngest fully expanded leaves of the main tiller (Wu *et al*., 2014), after 7 and 13 days of treatment. Plant material was harvested fifteen days after the onset of the experiment. The reduction of shoot and root growth were calculated as relative shoot/root dry weight (DW) = (shoot/root DW in treatment) / (shoot/root DW in the control). The concentration of Fe in shoots was quantified.

### QTL mapping and statistical analysis

Chromosome segment substitution lines (CSSL) maps were constructed using SSR markers, and SNP-markers from 6K Infinium platform and GBS platform (Arbelaez *et al*., 2015). Stepwise regression was used to select the most important segments for the traits evaluated and the likelihood ratio based on linear regression was used to estimate LOD scores. A permutation test using 1,000 permutations was carried out to determine the experiment-wise significance threshold (p = 0.05) and QTL detection was completed using RSTEP-LRT-ADD method in CSL program of QTL IciMapping v. 4.1 (Meng *et al*., 2015), according to (Arbelaez *et al*., 2015).

### Gas exchange measurements

Gas exchange was measured with a portable photosynthetic gas exchange system Li-Cor 6400-XT (LI-COR, Inc., Lincoln, NE, USA) 8 and 13 days after the onset of the treatment. The youngest fully-expanded leaf on the main tiller of each plant was evaluated between 10 am to 4 pm. Reference CO_2_ concentration, flow rate and PAR were set up at 400 μmol mol^-1^, 300 mmol s^-1^ and 1000 µmol m^-2^ s^-1^, respectively. Two plants of each genotype per replicate (n = 8) were used for the measurements in both treatments.

### Leaf spectral reflectance measurement

Leaf spectral reflectance was measured on the first and second fully-expanded leaves (6 plants per container; 4 containers) 7 and 14 days after the onset of the treatments. Normalized difference vegetation index (NDVI) was calculated as: NDVI = (R_NIR_ –R_RED_)/(R_NIR_ + R_RED_), where R represents the reflectance at a given wavelength (Huang *et al*., 2013). Renormalized difference vegetation index (RDVI) was calculated as: 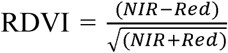 (Roujean and Breon, 1995). The carotenoid reflectance index (CRI) I and II were determined in order to estimate carotenoid concentration relative to chlorophyll concentration and were calculated as 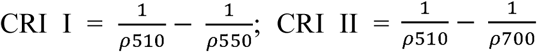 (Gitelson *et al*., 2002).

### LBS measurement

Leaf bronzing symptoms were scored on the two youngest fully expanded leaves of the main tiller, on a scale from 0 (no symptoms) to 10 (dead leaf) (Wu *et al*., 2014). The scoring was performed after 7 and 14 days of treatment.

### ROS staining in leaves

In-situ detection of ROS was conducted in the youngest fully expanded leaf as described (Wu *et al*., 2017). ROS formation was visualized as brown precipitation, documented by a camera (EOS 1200D, Canon, Tokyo, Japan).

### Electrolyte leakage

To evaluate the cell membrane damage (Bajji *et al*., 2002), the youngest fully expanded leaf was cut into small pieces of approximately 2 cm, washed three times with distilled water and stored in tubes filled with 30 mL of milli-Q water. Electrical conductivity measurements were taken at the moment of the harvest (0 h - ECi), 3, 6, 18 and 24 hours after harvest (ECf). After 24 h, the samples were autoclaved and a final measurement was performed. The electrolyte leakage was calculated as: (ECf-ECi)/(ECt-ECi)×100.

### Malondialdehyde assay

Malondialdehyde (MDA) was quantified as 2-thiobarbituric-responsive substances (Höller *et al*., 2014). Absorbance was measured at 440, 532 and 600 nm with a microplate reader (Powerwave XS2, BioTek) as described in (Rakotoson *et al*., 2019). Calculations of MDA concentrations included background corrections.

### Extraction of the root plaque

The root plaque from fresh roots was extracted with cold dithionite-citrate-bicarbonate (Taylor and Crowder, 1983), with some modifications. Rice roots were detached, washed once with distilled water and immersed in a solution of 20 mL of 0.3 M Na_3_C_6_H_5_0_7_·2H_2_O, 2.5 mL of 1.0 M NaHCO_3_ and 1.5 g of Na_2_S_2_O_4_ at room temperature. The solution was collected and the roots washed 3 times with 25 mL of distilled water. The resulting solution was made to final volume (100 mL) with distilled water and the washed roots were prepared for Fe determination by Atomic Absorption Spectrometry (AAS, Perkin-ELMER 1100B, Überlingen, Germany). Root oxidizing power was calculated as the amount of Fe precipitated on the root surface per root DW.

### Determination of iron content in different tissues

For the determination of Fe content in root and shoot, as well as the compartmentalization of Fe in shoot, leaf blades (LB) were separated from the sheath and culm (here referred as culm). Tissues were dried at 60 °C for seven days. Dry samples were ground to fine powder followed by digestion with 65% HNO_3_ in a microwave pressure digestion system (MARS6, CEM GmbH, Kamp-Lintfort, Germany). After digestion, samples were diluted to a final volume of 20 mL with distilled water. Fe content was determined by AAS (Perkin-ELMER 1100B).

### RNA extraction and transcriptomic analysis

Total RNA was extracted with peqGOLD RNA kit (Peqlab; Erlangen, Germany) following the manufacturers’ instructions. The genomic DNA was removed with RNAase-free with DNAase. The integrity of RNA was tested on a bleach agarose gel (Aranda *et al*., 2012) and for purity using NanoDrop OneC (Thermo Fisher Scientific, Braunschweig, Germany).

For transcriptomic analysis, approximately 10 µg of total RNA was used for high-throughput cDNA sequencing by Illumina HiSeq™ 4000 technology (BGI Genomics Co., Ltd – Denmark). RNAs derived from three biological replicates were used to generate the libraries. Eighteen libraries were constructed, from samples harvested seven days after the onset of treatments (control and +Fe). The cDNA libraries were prepared according to Illumina’s protocols. After sequencing, read quality was checked and all low-quality reads (PHRED value < 20) were removed.

### Expression Analysis

Genomes and genomic annotation for both *O. sativa* and *O. meridionalis* were obtained from Ensembl plants (https://plants.ensembl.org/index.html). Libraries from *O. sativa* cv. Curinga were aligned against the *O. sativa* genome, libraries from *O. meridionalis* were aligned against the *O. meridionalis* genome and libraries from CM23 were aligned against an *in silico* inferred CM23 genome (described below). Alignments were done using STAR (Dobin *et al*., 2013) on default settings and feature counting was performed with the *SummarizeOverlaps* function on Union mode from the *GenomicAlignments* (Lawrence *et al*., 2013) package for R. Principal component analyses were performed to cluster samples based on read count for quality control purposes. Subsequent differential expression analysis were performed (+Fe *vs*. control) through the standard protocol from DESeq2 (Love *et al*., 2014) package for R. Genes for which *q-value* < 0.05 were considered differentially expressed (DEG). The R package fgsea (Korotkevich and Sukhov, 2016) was applied to find Gene Ontology terms induced or repressed by the +Fe treatment (number of permutations = 10^5, *q-value* < 0.05). For cross-species comparisons of DEG, we built a one-to-one correspondence table between *O. sativa* and *O. meridionalis* genes based on homology information from Ensembl plants database. When, for a given gene, the homology relationship between species was of the type “one-to-many”, we selected the one with highest sequence alignment identity. Intersection diagrams of DEGs in different species were done with the R package upSetR (Conway *et al*., 2017). The data is publicly available through the GEO database with accession number GSE151559.

### CM23 hybrid genome reconstruction

CM23 genome is a result of hybridization of wild parental *O. meridionalis* (≈ 6.18%) with *O. sativa*, with three major introgressions on chromosomes 7, 9 and 10. Based on the approximated coordinates of those introgressions originated from *O. meridionalis* (Arbelaez *et al*., 2015), we used sequence homology to model the CM23 genome and used transcriptomic data to support the model. Assuming the introgressed DNA has to be placed at a homologous genomic segment, we considered *S*_*N*_ and *E*_*N*_ to be the start and end points of a given introgression coordinate in a chromosome *N* and defined as *RefSeq* the sequence from *O. sativa* on chromosome *N* from *S*_*N*_ to *E*_*N*_. Then we used the *NucMer* script from MUMMER (Kurtz *et al*., 2004) to align the *RefSeq* against the entire chromosome *N* of *O. meridionalis* with set parameters *maxgap* = 500 and *mincluster* = 100. Afterward, we filtered out aligned blocks that were both shorter than 2000 bases and had less than 90% identity. This large alignment was used to infer the introgression sequence (*i*.*e*., the *O. meridionalis* sequence that is homologous to the *RefSeq*).

We used RNAseq libraries from CM23, *O. sativa* and *O. meridionalis* to support our introgression prediction. Each library was mapped against both *O. sativa* and *O. meridionalis* genomes using STAR (Dobin *et al*., 2013) on default settings. Next, for each annotated exon in each evaluated chromosome, we calculated its coverage on each mapping job using *bedtools* (Quinlan and Hall, 2010). Then, in order to visualize coverage profile over the exons, we plotted line graphs as follows. First, we ranked each exon by its start position coordinate in the chromosome, which we called indexed position. Then a coverage profile (CP) for each indexed position *i* was calculated as: 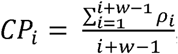, where ρ_i_ is the mean coverage of the exon whose indexed position is *i*, and w ∈ ℕ is an arbitrary window size that defines the graph line smoothness. We also defined dissimilarity distance *D* between two coverage profiles as D_i_ = (CP1_i_ - CP2*i*)^2^.

In summary, we artificially built the CM23 genome model by taking the *O. sativa* genome as a template and then proceeded to substitute genomic segments where introgressions occurred. Such segments were inferred according to homology and transcriptomic library coverage.

## Results

### Introgression lines show variation in Fe tolerance and accumulation

In order to identify Fe tolerant lines in the *O. sativa* x *O. meridionalis* population (hereafter CM), three-week-old plants were exposed to +Fe for 15 days. A hundred and eighteen phenotypic traits were evaluated, including biomass and growth-related traits, symptom formation, Fe concentration in shoot and spectral reflectance indexes of leaves (Table S1). The population showed significantly reduced shoot and root length under +Fe (Fig. S1A-B). When evaluating the relative DW, we observed that only eight (for shoots) and two lines (for roots) had significant decreases when submitted to +Fe, respectively (Fig. S1C-D). Symptoms related to stress, evaluated by LBS, began to develop after seven days of treatment. Thirteen days after the onset of the treatment *O. meridionalis* showed the lowest LBS (LBS = 1.0). Among the ILs, CM24 and CM23 were the most tolerant genotypes (1.25 and 2.25, respectively) (Fig. S1E).

Shoot Fe concentration and LBS were positively correlated in the CM population (Fig. 1). Considering all the lines, shoot Fe concentration explained only 11% (p = 0.05) of the observed variation in LBS. When we removed one outlier value (CM23 - Fe in shoot = 7.09 mg g^-1^ DW), the R^2^ value increased to 31.34% (p < 0.0007).

**Figure 1:**
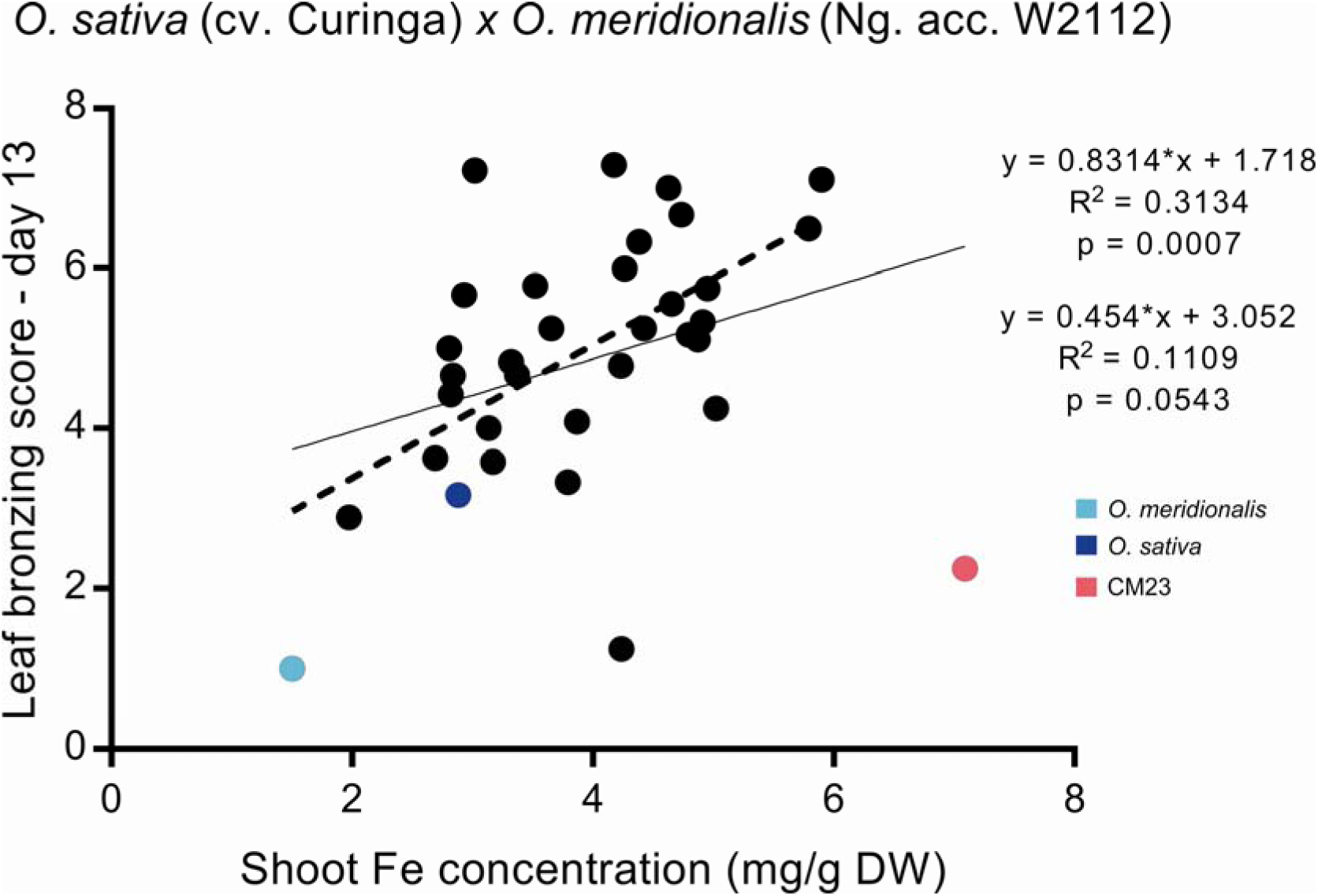
Linear regression of shoot Fe concentration *versus* leaf bronzing score (LBS) in *O. sativa* cv. Curinga x *O. meridionalis* (Ng. acc. W2112) (CM) population submitted to 250 mg L^-1^ of Fe for fifteen days. The dark blue data point represents the sensitive parental, *O. sativa* cv. Curinga. The light blue data point represents the tolerant parental *O. meridionalis* (Ng. acc. W2112). The pink data point represents the tolerant line with low symptom formation and high Fe concentration on shoots, named CM 23. The upper equation represents the values of correlation after the outlier value was removed.

### QTL analyses reveal regions associated with Fe tolerance

QTL analysis was performed to identify *loci* underlying the genetic variation of the evaluated phenotypic traits in the population. A stepwise regression analysis identified six different regions on chromosomes 1, 4 and 9 that were significantly associated with 15 different traits (LOD > 2.75) (Table 1). These putative QTLs explained from 16.5% (RDVI on day 7 – segment 1.1) up to 49% (RDVI on day 7 – segment 4.4) of phenotypic variability. The lines CM23 and CM24, which presented the lowest LBS while showing high Fe concentration in the shoot (Fig. 1), share a common introgression on chromosome 9, segment 9.4 (Arbelaez *et al*., 2015). This segment was associated with LBS and explained 42% of phenotypic variability (Table 1). The detection of putative QTLs for different traits in similar regions/segments (for example, segment 1.1, 4.4 and 9.4) suggests that these regions harbor genes involved in +Fe tolerance.

**Table 1:** Significant regions associated with traits evaluated during the screening of the population *O. sativa* cv. Curinga x *O. meridionalis* (Ng. acc. W2112) (LOD > 2.75) using a stepwise regression single marker analysis.

| Trait name | Chr | Segment | Marker position (bp) | LOD | PVE(%) | Add |
| --- | --- | --- | --- | --- | --- | --- |
| **OSAVId7** | 1 | 1.1 | 3088667 | 6.1 | 27.5 | 0.0355 |
| **TVId7** | 1 | 1.1 | 3088667 | 4.0 | 23.5 | 0.0475 |
| **Lic1d7** | 1 | 1.1 | 3088667 | 5.2 | 24.7 | 0.0299 |
| **SIPIId7** | 1 | 1.1 | 3088667 | 5.2 | 26.2 | 0.025 |
| **RDVId7** | 1 | 1.1 | 717702 | 3.5 | 16.6 | 0.0325 |
| **Gd5** | 4 | 4.4 | 18320014 | 2.8 | 34.1 | -0.0598 |
| **NDVId7** | 4 | 4.4 | 18320014 | 2.8 | 31.5 | -0.0483 |
| **OSAVId7** | 4 | 4.4 | 18320014 | 8.4 | 45.4 | -0.0518 |
| **TVId7** | 4 | 4.4 | 18320014 | 5.3 | 35.0 | -0.066 |
| **Lic1d7** | 4 | 4.4 | 18320014 | 7.7 | 44.9 | -0.0457 |
| **SIPIId7** | 4 | 4.4 | 18320014 | 7.1 | 41.5 | -0.0357 |
| **RDVId7** | 4 | 4.4 | 18320014 | 7.5 | 48.7 | -0.0558 |
| **CRI1d7** | 9 | 9.2 | 9195305 | 3.2 | 35.9 | -0.0507 |
| **Lic2d7** | 9 | 9.3 | 12154616 | 3.1 | 37.8 | -0.0715 |
| **LBS13** | 9 | 9.4 | 17948535 | 4.0 | 41.9 | 1.8188 |
| **Lic2d5** | 9 | 9.4 | 17948535 | 3.0 | 34.2 | -0.0625 |
| **PRIId7** | 9 | 9.4 | 17948535 | 3.0 | 33.6 | -0.1578 |
| **ZMIId13** | 9 | 9.4 | 19065062 | 2.7 | 31.7 | -0.0602 |
| **Lic2d13** | 9 | 9.4 | 19065062 | 3.5 | 39.5 | -0.0902 |
| **GMIId13** | 9 | 9.5 | 21802016 | 3.7 | 39.4 | -0.0763 |

Chr: chromosome; LOD: logarithm of odds; PVE: phenotypic variability explained; Add: additive effect; OSAVI: Optimized Soil-Adjusted Vegetation Index; TVI: Triangular Vegetation Index; Lic1 and Lic2: Lichtenthaler Indexes; SIPI: Structure Intensive Pigment Index; RDVI: Renormalized Difference Vegetation Index; G: Greenness Index; NDVI: Normalized Difference Vegetation Index; CRI1: Carotenoid Reflectance Index 1; LBS: leaf bronzing score; PRI: Photochemical Reflectance Index; ZMI: Zarco-Tejada & Miller Index; GMI: Gitelson and Merzlyak Index; d indicates the day after the start of the treatment.

Based on the screening and QTL analysis, CM23 was the most tolerant line from the CM population despite very high shoot Fe concentration (Fig. 1). This line and the parental genotypes, *O. sativa* and *O. meridionalis*, were used in subsequent experiments for physiological, biochemical and molecular analyses.

### Introgression line CM23 and *O. meridionalis* are more tolerant to high Fe than *O. sativa*

Based on LBS data *O. meridionalis* and CM23 were more tolerant to Fe excess than *O. sativa* after seven and fourteen days of treatment (Fig. 2A). All three genotypes showed similar growth reduction under +Fe compared to the plants in the control (Fig. 2B-E). CM23 showed less reduction compared with *O. sativa* (Fig. 2C-E).

**Figure 2:**
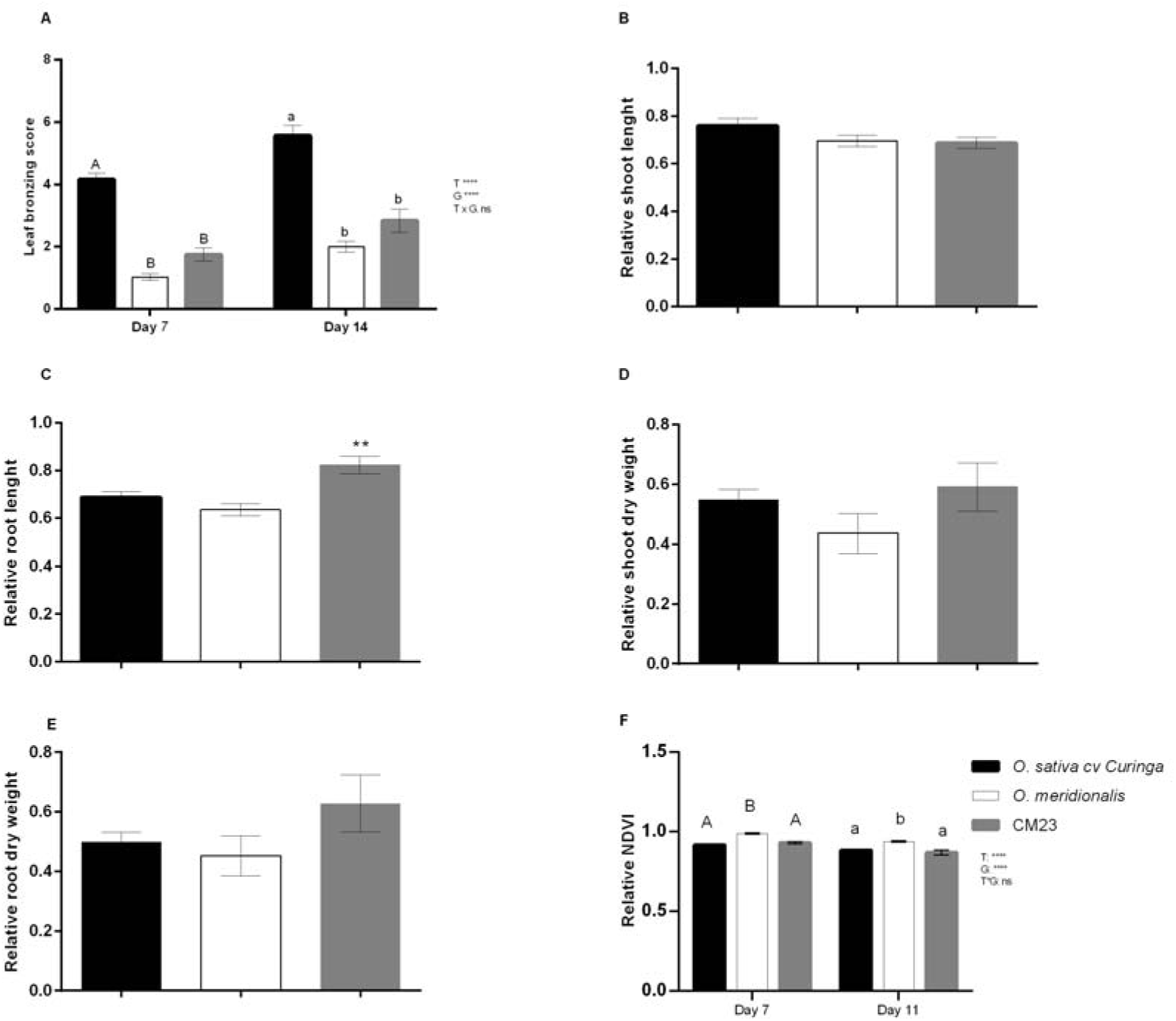
Responses of parents and selected introgression lines exposed to Fe toxicity (250 mg L^-1^ Fe^2+^) for 14 days. (A) Leaf bronzing score in Fe treatment after seven and fourteen days of treatment (n = 16). (B) Relative shoot length (n = 16). (C) Relative root length (n = 16). (D) Relative shoot dry weight (n = 16). (E) Relative root dry weight (n = 16). Values are the averages ± SE. Asterisks indicate statistical difference when compared with *O. sativa* (Student *t*-test, **P-value < 0.01). Different letters above the bars indicate the differences were significant at *P* < 0.05 by *post hoc* Tukey’s test. Uppercase represents comparisons in seven days and lowercase comparisons between fourteen days after onset of Fe toxicity. (F) Relative normalized difference vegetation index (NDVI) on days 7 and 11 after the onset of treatment (n = 16). Different letters above the bars indicate the differences were significant at *P* < 0.05 by *post hoc* Tukey’s test. Uppercase represents comparisons between genotypes after seven days and lowercase comparisons between genotypes after eleven days of Fe excess treatment.

Leaf spectral reflectance indexes were used to monitor stress. The wild parental had higher relative values of NDVI than *O. sativa* and CM23 (Fig. 2F). Carotenoids are associated with light absorption in plants, and protect them from photo-oxidation. Stressed plants contain higher concentrations of carotenoids (Gitelson *et al*., 2002). Consistently, Fe treatment significantly increased the CRI I and II in *O. meridionalis* after seven days of treatment (Fig. S2A-B). However, the relative values detected in CM23 for these same parameters were always below/near to 1, indicating lower concentrations of carotenoids accumulated in this IL (Fig. S2A-B).

Changes caused by +Fe on photosynthesis in all the three lines were investigated by gas exchange analysis. We found that the net photosynthetic rate in all three genotypes was clearly affected by Fe treatment, with negative values in CM23 starting after eight days under +Fe (Fig. S3A-B). Stomatal conductance and leaf transpiration rate were also significantly affected by the Fe treatment in all three genotypes (Fig. S3C-D). However, the relative values of CM23 were significantly higher than in both parents, indicating that it maintains higher transpiration and conductance under +Fe.

### CM23 and *O. meridionalis* show lower oxidative stress under Fe stress

Seven days after the onset of the treatment, MDA concentration significantly increased in all genotypes when submitted to +Fe compared to control (Fig. 3A). *O. meridionalis* showed lower MDA concentration under +Fe compared to *O. sativa* and CM23 (Fig. 3A). After 14 days of +Fe, MDA increased in all genotypes compared to controls (Fig. 3B). However, values were significantly higher in *O. sativa* compared to *O. meridionalis* and CM23. CM23 also showed the lowest electrolyte leakage value when compared with both parents, corroborating the tolerant character of this line (Fig. 3C).

**Figure 3:**
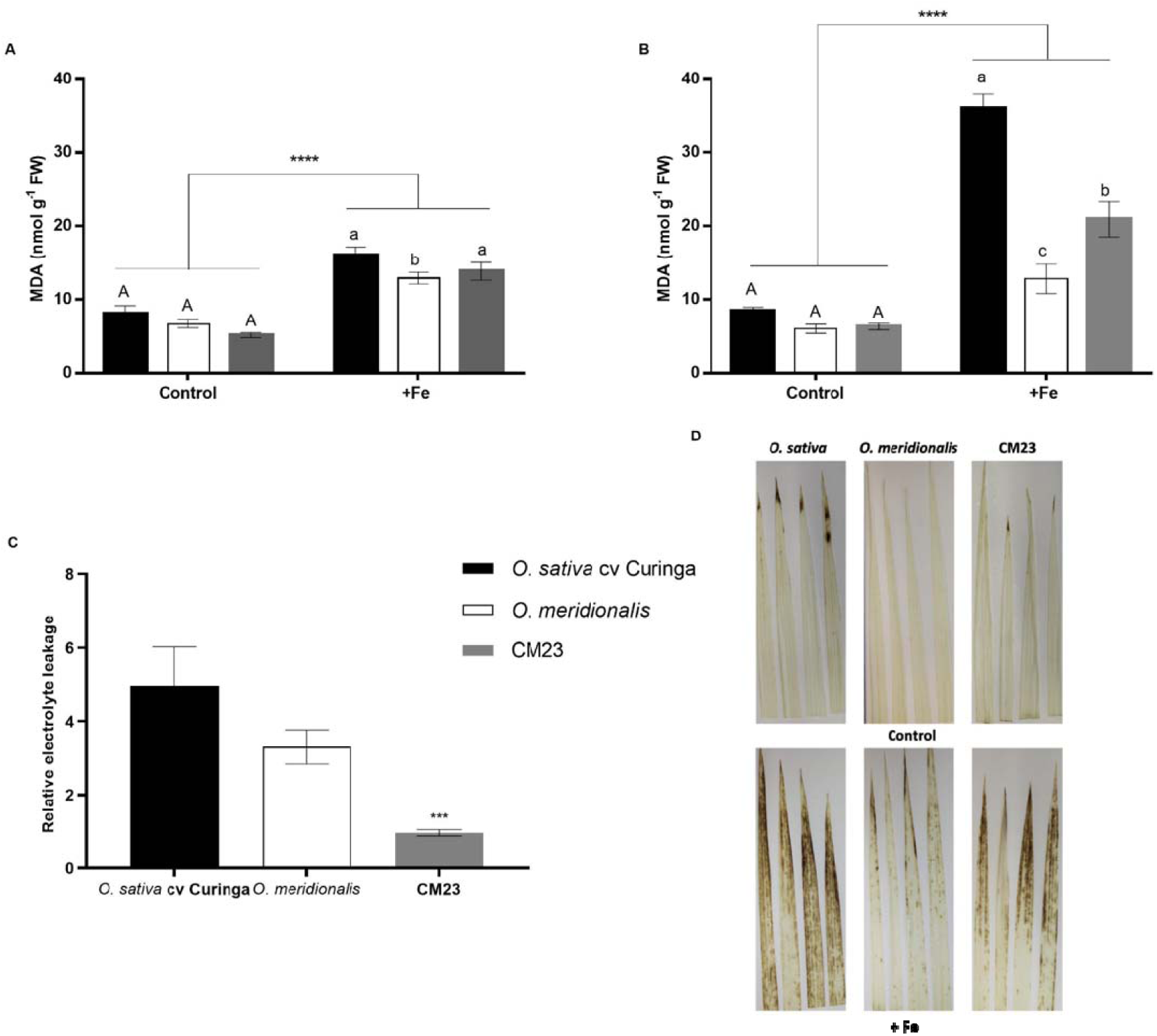
Physiological analysis of the tolerance to Fe toxicity. Shoot MDA concentration in contrasting parents (*O. sativa* cv. Curinga and *O. meridionalis* (Ng. acc. W2112) and in IL CM23 submitted to control and +Fe (250 mg L^-1^ Fe^2+^). Plants were submitted to +Fe for seven (A) and fourteen (B) days. (n = 4). Values are the averages ± SE. Asterisks indicate statistical difference between control and +Fe (Student *t*-test, ***P-value < 0.001). Different letters above the bars indicate the differences were significant at *P* < 0.05 by *post hoc* Tukey’s test. Uppercase represents comparisons between genotypes in control and lowercase comparisons in Fe toxicity. (C) Relative electrolyte leakage in leaves of parents and selected introgression line (CM23) submitted to control condition and Fe toxicity for 14 days (n = 16). (Student *t*-test, ***P-value < 0.001). (D) Hydrogen peroxide staining using 3,3′-Diaminobenzidine of parents and selected introgression line (CM23) under control and Fe toxicity (250 mg L^-1^ Fe^2+^ for 7 days) (n=4).

DAB staining was employed to visualize H_2_O_2_ generation in leaves (Fig. 3D). In the control, no apparent differences in H_2_O_2_ staining were observed in *O. meridionalis, O. sativa* and CM23. With the exception of *O. meridionalis*, all genotypes had a small brown stains at the tip of the leaves, even in control. However, when plants were cultivated under +Fe, brown precipitates indicating H_2_O_2_ production were observed in the leaves of *O. sativa* (Fig. 3D), whereas lower levels were found in CM23, and even less was observed in *O. meridionalis*. These results indicate the high level of tolerance to Fe excess of *O. meridionalis* and CM23.

### CM23 stores Fe in leaf sheaths and *O. meridionalis* excludes Fe from shoots

Fe concentration was analyzed in three different plant organs: LB, culm + sheath and roots. Fe treatment significantly increased Fe concentration in all shoot organs, as expected (Fig. 4A-B). No genotypic differences in Fe concentration in culm and sheath were observed under control. However, under +Fe, the highest Fe concentration in culm + sheath was observed in CM23 and lowest Fe concentration in the wild parent (Fig. 4A). The Fe concentration in the LB did not show any genotypic variation under control or +Fe treatment (Fig. 4B).

**Figure 4:**
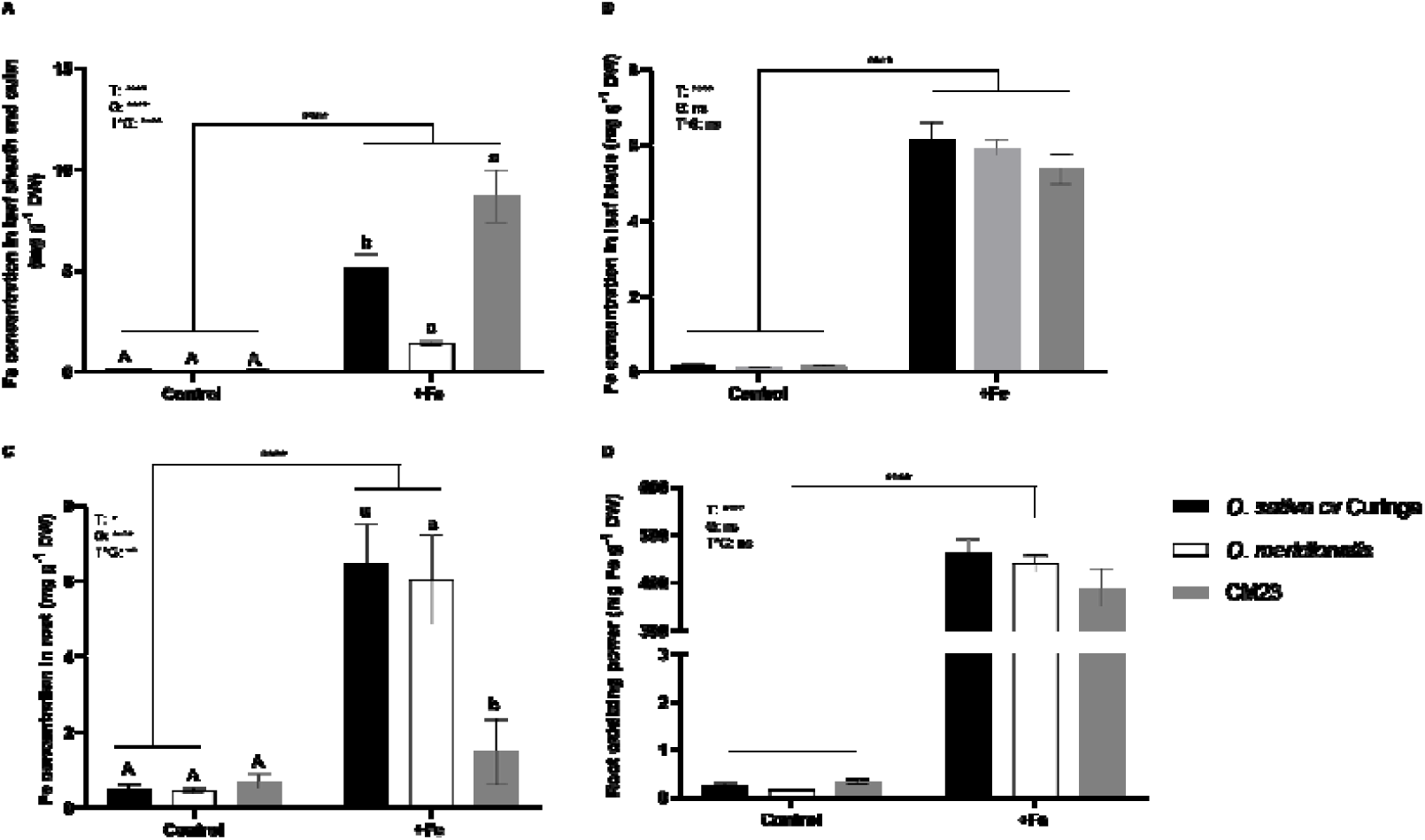
Fe concentration in three different organs of *O. sativa* cv. Curinga, *O. meridionalis* (Ng. acc. W2112) and in CM23 exposed to control conditions and to Fe toxicity (250 mg L^-1^ Fe^2+^) for fourteen days. (A) Fe concentration in leaf sheath and culm (mg g^-1^ DW). (B) Fe concentration in leaf blades (mg g^-1^ DW). (C) Fe concentration in roots (mg g^-1^ DW). (D) Root oxidation power (Fe concentration in the iron plaque) (mg Fe g^-1^ DW). (n = 8). Values are the averages ± SE. Different letters above the bars indicate that the differences were significant at *P* < 0.05 by *post hoc* Tukey’s test. Uppercase letters represent comparisons in control and lowercase comparisons between genotypes under Fe toxicity. Asterisks indicate statistical difference between plants grown under CC and +Fe conditions (Student *t*-test, ****P-value < 0.0001).

Only *O. sativa* and *O. meridionalis* responded with increased root Fe concentration under +Fe when compared with plants cultivated under control (Fig. 4C). CM23 had the lowest Fe concentration in roots, suggesting that the Fe absorbed by CM23 is translocated to shoot tissues. When the root oxidizing power was evaluated, only treatment differences were observed (Fig. 4D), suggesting that the tolerance presented by CM23 is based on a shoot mechanism.

### Transcriptomic analysis of CM23 and parental lines under control and +Fe

Transcriptomic analyses were performed on shoots of *O. sativa, O. meridionalis* and CM23 plants exposed to control and +Fe conditions. A total of 4,171 genes (1,578 down and 2,593 up-regulated) were differentially expressed in *O. sativa*, accounting for 10.73% of all annotated genes (Table S2 and Fig. 5). In *O. meridionalis*, a total of 5,601 DEG (2,739 down and 2,862 up-regulated - Table S3 and Fig. 5) were identified (18.52 % of all annotated genes), whereas CM showed 1,449 (643 down and 806 up-regulated, with 27 and 47 introgressed genes, respectively) were DEG (3.78 % of all annotated genes) (Table S4 and Fig. 5).

**Figure 5:**
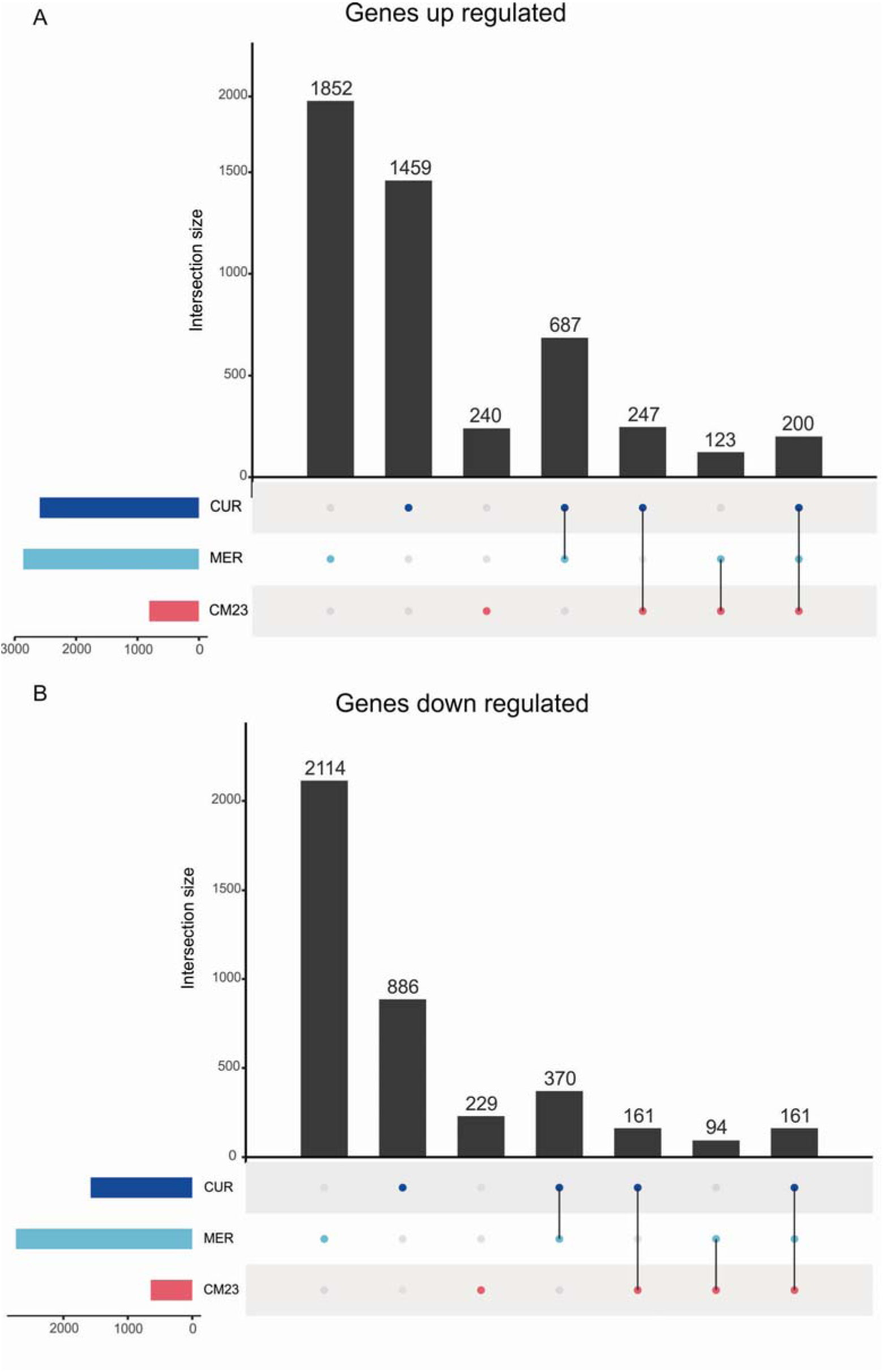
Upset chart with genes up and down-regulated by Fe toxicity (250 mg L^-1^ Fe^2+^) seven days after + Fe treatment in *O. sativa* cv. Curinga, *O. meridionalis* (Ng. acc. W2112) and the IL CM23. (A) Genes up and (B) down-regulated by Fe treatment in *O. meridionalis, O. sativa* cv. Curinga, CM23, genes commonly regulated between *O. sativa* and *O. meridionalis, O. sativa* and CM23, *O. meridionalis* and CM23, and commonly regulated in all three genotypes. CUR: *O. sativa* cv. Curinga; MER: *O. meridionalis;* CM23: IL CM23.

When considering genes exclusively regulated in one genotype, we found 1,459, 1,852 and 240 exclusively up-regulated in *O. sativa, O. meridionalis* and CM23, respectively, and 886, 2,114 and 229 genes exclusively down-regulated (Fig. 5A-B and Tables S5 and S6). After the inference of homologies between *O. sativa* and *O. meridionalis*, we identified genes commonly regulated in different genotypes. A total of 687 and 370 DEG were up and down-regulated both in *O. sativa* and *O. meridionalis* respectively, 247 and 161 DEG up and down-regulated both in *O. sativa* and the IL CM23 respectively, and 123 and 94 DEG up and down-regulated both in *O. meridionalis* and CM23 respectively. However, when considering all genotypes, a total of 200 and 161 genes were up and down-regulated in common (Fig. 5A-B).

### Gene set enrichment analysis (GSEA) of transcripts associated with +Fe response indicates similar processes regulated in all genotypes

Aiming to compare the transcriptomic changes in shoots under +Fe in *O. sativa, O. meridionalis* and CM23, we performed GSEA. We found 38 GO terms enriched in *O. sativa* (18 up and 20 down-regulated by +Fe), 31 GO terms enriched in *O. meridionalis* (16 up and 15 down-regulated) and 28 GO terms enriched in CM23 (16 up and 12 down-regulated) (Fig. 6 and 7).

**Figure 6:**
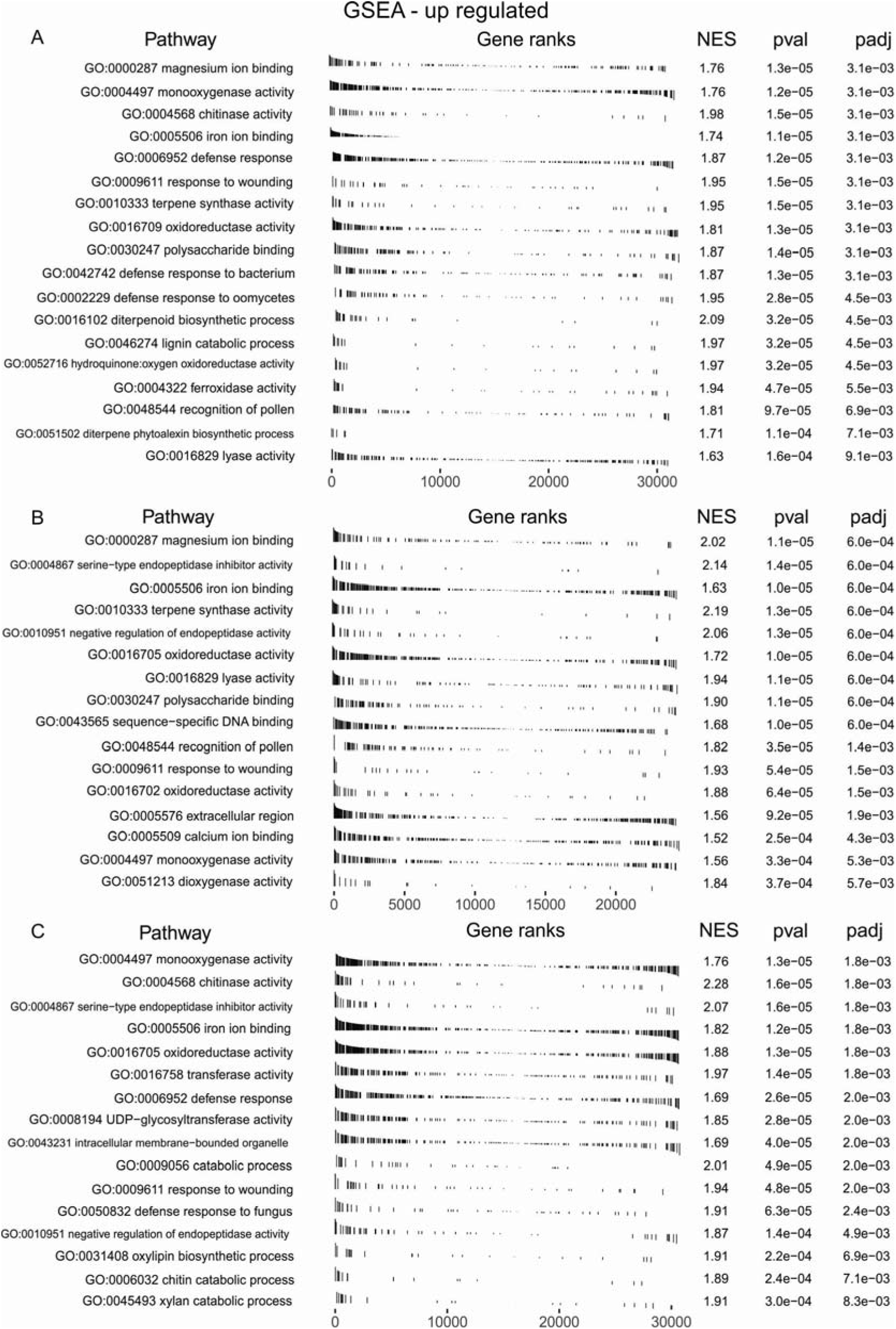
Gene set enrichment analysis of up-regulated genes in the transcriptome of shoots from *O. sativa* cv. Curinga, *O. meridionalis* (Ng. acc. W2112) and IL CM23 seven days after the onset of treatment with Fe excess (250 mg L^-1^ Fe^2+^). The line represents the genes ordered by logFC. The higher the logFC of a gene, the more to left the line corresponding to that gene is represented.

**Figure 7:**
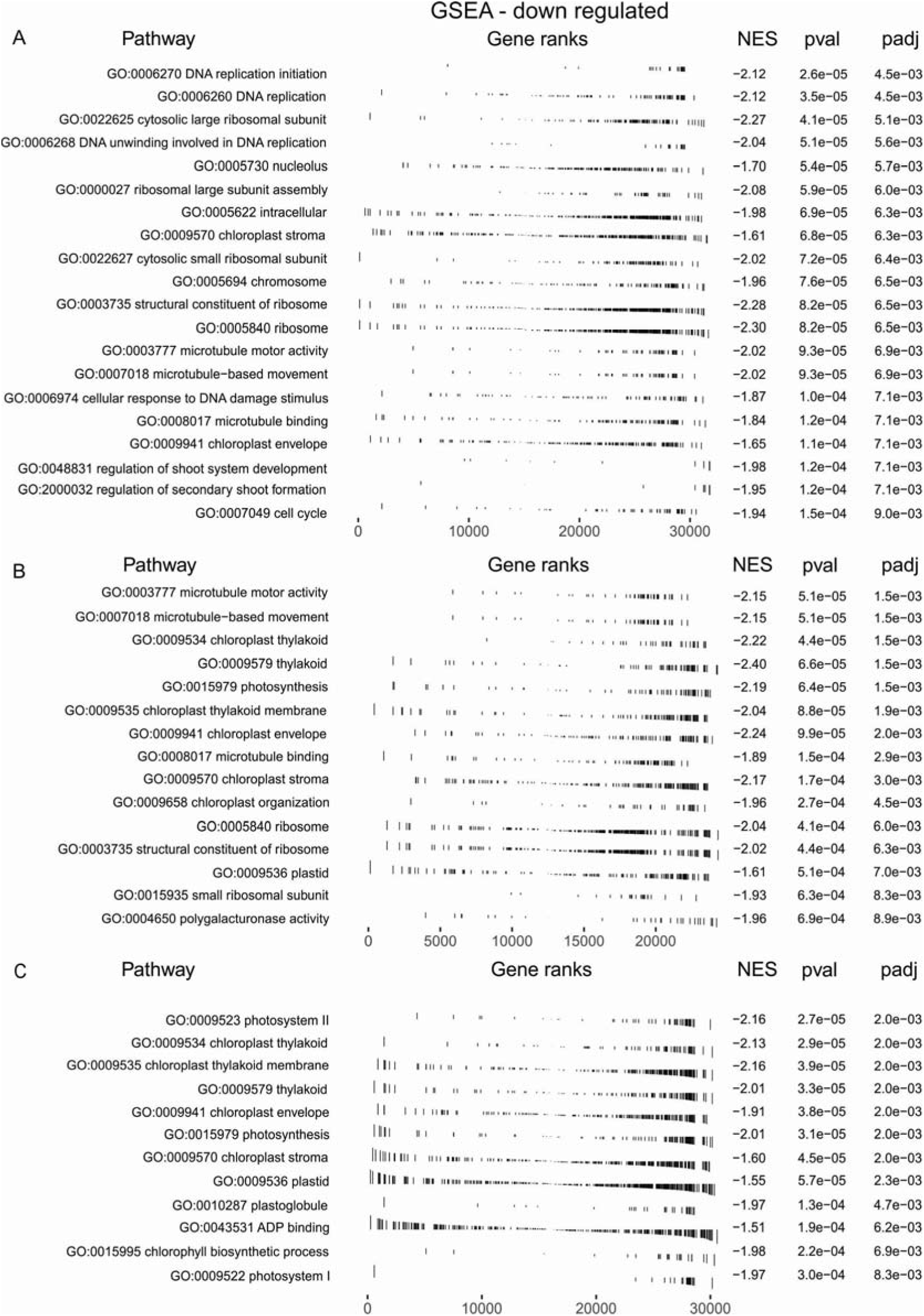
Gene set enrichment analysis of down-regulated genes on the transcriptome from shoots of *O. sativa* cv. Curinga, *O. meridionalis* (Ng. acc. W2112) and IL CM23 seven days after the onset of treatment with Fe excess (250 mg L^-1^ Fe^2+^). The line represents the genes ordered by logFC. The higher the logFC of a gene, the more to right the line corresponding to that gene is represented.

Only three terms were overrepresented among up-regulated genes in all genotypes: “iron ion binding”, “monooxygenase activity” and “response to wounding” (Fig. 6A-C). On the other hand, two terms were overrepresented among down-regulated genes in all genotypes and both were associated with chloroplast: “chloroplast stroma” and “chloroplast envelope” (Fig. 7A-C).

Five terms were identified as equally overrepresented and up-regulated in *O. sativa* and *O. meridionalis*: “magnesium ion binding”, “terpene synthase activity”, “polysaccharide binding”, “recognition of pollen” and “lyase activity” (Fig. 6A-B). Two GO terms were commonly up-regulated between *O. sativa* and CM23: “chitinase activity” and “defense response”, suggesting a biotic stress response. *O. sativa* and CM23 did not share overrepresented GO terms for down-regulated genes.

Even though the number of GO terms up-regulated in CM23 was lower than in the parental genotypes, some of them were exclusively overrepresented: “transferase activity, transferring hexosyl groups”, “UDP−glycosyltransferase activity”, and “intracellular membrane−bounded organelle” (Fig. 6C). On the other hand, the down-regulated terms exclusively overrepresented in CM23 followed the same pattern as those overrepresented the *O. meridionalis* (“photosystem II”, “photosystem I”, “plastoglobule”, “chlorophyll biosynthetic process” and “ADP binding”) (Fig. 7C).

### Fe responsive genes in the tolerant CM23

In order to identify possible pathways and mechanisms linked to Fe tolerance in the interspecific hybrid, we focused on genes exclusively regulated in CM23 in comparison to the sensitive *O. sativa*. In contrast, we considered genes that were equally responsive to Fe stress in both the *O. sativa* parent and CM23 with a logFC >2 (Table S7) as stress response rather than stress tolerance genes.

We found that 377 genes were exclusively up-regulated in the tolerant CM23 (Table S8). These included a number of genes that could be involved in Fe homeostasis and detoxification, especially two genes encoding metallothionein genes (Os01g0200700, *OsMTI-3a* and Os05g0202800, *OsMTI-3b*). Another candidate would be an iron-responsive element-binding gene (Os08g0191100), which catalyzes the isomerization of citrate to isocitrate via cis-aconitate. Five transcription factors (TF) from the bHLH (basic helix-loop-helix) family, which has members involved in the regulation of Fe homeostasis (Gao *et al*., 2019), were exclusively induced by +Fe in CM23. Also, a variety of transporters was contained in this list of genes, including potassium transporter OsHAK9 (Os07g0679000), high-affinity nitrate transporter OsNar2.2 (Os04g0480200), molybdate transporter OsMOT1 (Os08g0101500), copper transporter OsCOPT3 (Os03g0370800), and the sugar transport protein OsMST2 (Os03g0594400), which mediates active uptake of hexoses by sugar:proton symport. The heavy metal transport/detoxification protein (Os01g0976300) constitutes another plausible candidate that could be involved in Fe tolerance. Eight peroxidases were identified as induced by +Fe treatment in CM23 (Table S8), and could be involved in the removal of H_2_O_2_, oxidation of toxic reductants and general response to oxidative stress. We also identified several genes associated with osmotic stress and tolerance to salt and drought such as *Os-NADP-ME2* (Os01g0723400), *OsTIFY11a* (Os03g0180800), *OsMRS2* (Os01g0955100) and *OsMBL1* (Os01g0348900). These results suggest that osmoregulation could play a role in the tolerance to +Fe observed in CM23.

We further found that 332 genes exclusively down-regulated in CM23 compared to *O. sativa*. Among those were five TFs from the bHLH family and six TFs from the zinc finger family, as well as a gene from the WRKY family (*OsWRKY57 -* Os12g0102300). Another category of downregulated genes in CM23 was transporters, including two genes from the oligopeptide transporter family, *OsOPT8* (Os02g0695800) and *OsOPT3* (Os06g0125500), one gene encoding a peptide transporter PTR2 (Os06g0239500), two genes from the ABC family transporters *OsABCG2* (Os01g0615500) and *OsABCC7* (Os04g0588700), and *OsNRAMP4* (Os01g0503400) (Table S8). In agreement with the data obtained from GSEA analysis, a great number of genes associated with photosynthesis such as genes coding for chlorophyll a/b-binding protein (Os02g0197600), ribulose bisphosphate carboxylase/oxygenase activase (Os04g0658300) and photosystem II stability/assembly factor HCF136 (Os06g0729650) were down-regulated under +Fe (Table S8). The same expression patterns was seen for the stay green-like protein (Os04g0692600), which promotes chlorophyll degradation in leaves (Rong *et al*., 2013) was also down-regulated in CM23 by +Fe.

### Fe-responsive genes introgressed from *O. meridionalis* into CM23

We further focused on genes that were both Fe-responsive and also located within the introgression segments of CM23. Out of 47 DEG introgressed from *O. meridionalis* and up-regulated by +Fe treatment, 20, 15 and 12 were located in the introgressions on chromosomes 7, 9 and 10, respectively. For eight out of these 47 genes, it was not possible to infer a homologue in *O. sativa*. From these eight *O. meridionalis*-specific genes, three are located on chromosome 7, three on chromosome 9 and two genes on chromosome 10 (Table 2).

**Table 2:** Differentially expressed genes introgressed from *O. meridionalis* into CM23.

| Gene_ID | logFC | q value | Homologous gene | Description homologous |
| --- | --- | --- | --- | --- |
| Up-regulated genes |  |  |  |  |
| OMERI07G00330 | 1.065 | 0.0097 | Os07g0114000 | Similar to Protein phosphatase 2C-like |
| OMERI07G00620 | 1.848 | 0.005 | - | - |
| OMERI07G01050 | 1.588 | 0.032 | Os07g0127500 | Similar to PR-1a pathogenesis related protein (Hv-1a) precursor |
| OMERI07G01070 | 1.647 | 0.020 | - | - |
| OMERI07G01130 | 3.062 | 6.72E-15 | - | - |
| OMERI07G01900 | 0.733 | 0.044 | Os07g0151200 | Sugar/inositol transporter domain containing protein |
| OMERI07G02190 | 1.114 | 0.00014 | Os07g0158800 | Protein of unknown function DUF266, plant family protein |
| OMERI07G02920 | 1.530 | 0.008 | Os07g0168300 | Similar to Glutathione S-transferase GSTU6 |
| OMERI07G02960 | 1.611 | 3.23E-5 | Os07g0170000 | Similar to Brn1-like protein |
| OMERI07G03280 | 1.219 | 0.011 | Os07g0174400 | Similar to Non-specific lipid-transfer protein |
| OMERI07G03370 | 1.462 | 0.002 | Os07g0175600 | Plant lipid transfer protein and hydrophobic protein, helical domain containing protein |
| OMERI07G04020 | 1.141 | 0.002 | Os07g0192000 | ATPase, AAA-type, core domain containing protein |
| OMERI07G05270 | 1.682 | 8.07E-9 | Os07g0201800 | UDP-glucuronosyl/UDP-glucosyltransferase domain containing protein |
| OMERI07G05590 | 0.716 | 0.005 | Os07g0206600 | Similar to Hexose transporter |
| OMERI07G06160 | 4.076 | 3.51E-12 | Os07G0218700 | Cytochrome P450 family protein |
| OMERI07G06460 | 1.383 | 0.005 | Os07g0231800 | Conserved hypothetical protein |
| OMERI07G06790 | 2.340 | 0.002 | Os08g0412700 | Protein of unknown function DUF1262 family protein |
| OMERI07G06870 | 1.404 | 0.0001 | Os07g0241800 | UDP-glucuronosyl/UDP-glucosyltransferase family protein |
| OMERI07G07720 | 0.874 | 0.039 | Os10g0566400 | Similar to RING-H2 finger protein ATL3J |
| OMERI09G04710 | 2.031 | 0.021 | Os09g0333000 | ABC transporter-like domain containing protein |
| OMERI09G05070 | 2.420 | 7.369E-6 | Os09g0344500 | Similar to O-methyltransferase ZRP4 (EC 2.1.1.) |
| OMERI09G06280 | 1.064 | 0.001 | Os09g0370500 | VQ domain containing protein |
| OMERI09G06320 | 1.162 | 0.012 | Os09g0371300 | Similar to predicted protein |
| OMERI09G06770 | 1.465 | 0.001 | Os09g0379600 | Similar to Homeobox-leucine zipper protein HOX25 |
| OMERI09G06970 | 1.368 | 0.001 | Os09g0385700 | Zinc finger, AN1-type domain containing protein |
| OMERI09G07290 | 3.897 | 1.869E-7 | - | - |
| OMERI09G07310 | 2.636 | 0.015 | - | - |
| OMERI09G08230 | 1.155 | 0.005 | Os09g0418000 | Similar to CBL-interacting protein kinase 16 |
| OMERI09G08730 | 1.167 | 0.017 | Os09g0431300 | Similar to MYB transcription factor MYB71 (Fragment) |
| OMERI09G09130 | 1.165 | 0.003 | Os09g0439200 | Jasmonate ZIM-domain protein, Jasmonate-induced resistance to bacterial blight, Repressor of jasmonic acid signalin |
| OMERI09G10000 | 1.856 | 0.007 | Os09g0454600 | Similar to Mitochondrial phosphate transporter (Fragment) |
| OMERI09G10200 | 2.953 | 0.033 | Os09g0457900 | AP2/ERF transcription factor, Regulation of the internode elongation |
| OMERI09G10280 | 1.385 | 0.028 | Os08g0478466 | Protein of unknown function DUF296 domain containing protein |
| OMERI09G10540 | 1.379 | 0.003 |  |  |
| OMERI10G12860 | 1.116 | 0.017 | Os10g0524300 | LysM domain-containing protein, Embryo sac development |
| OMERI10G13020 | 2.167 | 1.33E-7 | Os10g0527601 | Glutathione S-transferase, C-terminal-like domain containing protein |
| OMERI10G13050 | 0.750 | 0.006 | Os10g0529400 | Similar to Glutathione S-transferase GSTU6 |
| OMERI10G13130 | 1.584 | 3.86E-7 | Os10g0530600 | Similar to Glutathione S-transferase GST 20 (EC 2.5.1.18) |
| OMERI10G13260 | 1.337 | 0.027 | Os10g0532300 | Heavy metal transport/detoxification protein domain containing protein |
| OMERI10G13550 | 1.873 | 0.005 |  |  |
| OMERI10G13880 | 1.390 | 0.0004 | Os10g0542800 | Brassinosteroid-signaling kinase, A member of the receptor-like cytoplasmic kinase (RLCK)-XII sub group, Major regulator in rice immunity |
| OMERI10G14200 | 2.331 | 0.006 | Os10g0548100 | Similar to DM280 protein |
| OMERI10G14220 | 1.058 | 0.011 | Os10g0548400 | Similar to F25C20.9 |
| OMERI10G14500 | 1.533 | 0.012 |  |  |
| OMERI10G14970 | 1.739 | 0.001 | Os10g0580400 | High-affinity urea transporter, Effective urea acquisition and utilisation, Effective use of low external urea as a N source |
| OMERI10G15240 | 1.098 | 6.36E-5 | Os10g0576600 | Tetratricopeptide-like helical domain containing protein |
| **Down-regulated genes** |  |  |  |  |
| OMERI07G00150 | -1.192 | 0.0002 | Os07g0108300 | Similar to Alanine aminotransferase |
| OMERI07G00410 | -0.864 | 0.026 | Os07g0115300 | Similar to Peroxidase2 precursor (EC 1.11.1.7) |
| OMERI07G01650 | -0.825 | 0.030 |  |  |
| OMERI07G02060 | -0.851 | 0.006 | Os07g0152900 | Similar to Glycolate oxidase (EC 1.1.3.15) (Fragment) |
| OMERI07G02290 | -1.188 | 0.0009 | Os07g0159700 | Protein kinase, catalytic domain domain containing protein |
| OMERI07G02340 | -0.857 | 0.035 | Os07g0160400 | Glyoxalase/bleomycin resistance protein/dioxygenase domain containing protein |
| OMERI07G03430 | -0.660 | 0.048 | Os07g0176900 | Similar to Ribose-5-phosphate isomerase precursor (EC 5.3.1.6) |
| OMERI07G03910 | -1.011 | 0.005 | Os07g0191200 | Plasma membrane H<sup>+</sup> ATPase (EC 3.6.3.6) |
| OMERI07G05950 | -1.080 | 0.0005 | Os07g0212200 | Similar to mRNA-binding protein (Fragment) |
| OMERI09G03920 | -0.694 | 0.020 | Os09g0307300 | Targeting for Xklp2 family protein |
| OMERI09G04970 | -1.204 | 0.010 | Os09g0343200 | Ankyrin repeat containing protein |
| OMERI09G05320 | -4.16 | 1.48E-5 | Os08g0327200 | OsPGL4 |
| OMERI09G05460 | -1.982 | 6.77E-6 | Os09g0350900 | Protein kinase, core domain containing protein |
| OMERI09G05480 | -4.369 | 6.39E-7 |  |  |
| OMERI09G05570 | -2.252 | 5.49E-6 |  |  |
| OMERI09G06420 | -1.704 | 0.017 |  |  |
| OMERI09G06460 | -1.185 | 0.023 |  |  |
| OMERI09G07520 | -0.860 | 0.015 | Os09g0402300 | Phosphatidylinositol-4-phosphate 5-kinase family protein |
| OMERI09G08870 | -1.169 | 2.16E-5 | Os09g0434500 | Similar to Ethylene response factor 2 |
| OMERI09G09840 | -0.850 | 0.040 | Os09g0452800 | Peptidase A1 domain containing protein |
| OMERI09G10700 | -0.904 | 0.013 |  |  |
| OMERI09G10710 | -0.787 | 0.019 | Os09g0477900 | NusB/RsmB/TIM44 domain containing protein |
| OMERI10G13120 | -1.279 | 0.0005 | Os10g0530500 | Similar to Glutathione-S-transferase Cla47 |
| OMERI10G13320 | -0.824 | 0.024 | Os10g0533500 | Similar to Beta-ring hydroxylase (Fragment) |
| OMERI10G13640 | -0.917 | 0.028 | Os10g0539900 | Tonoplast monosaccharide transporter, Vacuolar sugar transporter |
| OMERI10G14830 | -0.923 | 0.003 |  |  |
| OMERI10G15060 | -1.785 | 0.0004 | Os10g0578950 | Similar to 4-coumarate--CoA ligase-like 2 |

Four genes similar to Glutathione S-transferase (GST) could play an important role in Fe tolerance, as GSTs are multifunctional enzymes contributing to cellular detoxification. Additional interesting genes that could be involved in Fe tolerance include one sugar/inositol transporter (OMERI07G01900/Os07g0151200); one putative hexose transporter (OMERI07G05590/Os07g0206600) that belongs to the major facilitator superfamily (MFS); one heavy metal transport/detoxification (OMERI10G13260); and the transporter urea-proton symporter DUR3 (OMERI10G14970) that plays an important role in high-affinity urea uptake by cells at low external urea concentrations (Wang *et al*., 2012).

Genes encoding TFs from four different families were up-regulated (Table 2). The homeobox-leucine zipper protein HOX25 (OMERI09G06770) belongs to the HD-Zip I gene family of TF. The gene *OsSAP17* (OMERI09G06970 - zinc finger, AN1-type) encodes a protein with the A20/AN1 zinc-finger domain and plays a role in responses to abiotic stress (Vij and Tyagi, 2006). Other TFs identified are OMERI09G10200 which belongs to the APETALA2 (AP2)/Ethylene-Responsive Element Binding Factor family, and OMERI09G08730, similar to MYB TF family. Since TFs control the expression of numerous downstream genes providing stress tolerance to plants, these genes from *O. meridionalis* introgressed into CM23 genome could play essential roles in Fe tolerance.

From the introgression located on chromosome 9, we identified one ABC transporter-like domain containing protein - *ABCG47* (OMERI09G04710), and one gene coding for one acetylserotonin O-methyltransferase - *ASMT1* (OMERI09G05070) that is the final enzyme in a biosynthetic pathway that produces melatonin, which could provide protection against oxidative stress (Park *et al*., 2013). Also, one CBL-interacting protein kinase 16 (OMERI09G08230), which has GO terms related to sodium ion transport and hyperosmotic salinity response, was also identified as induced by +Fe.

On the other hand, a total of 27 genes introgressed from *O. meridionalis* were down-regulated by +Fe treatment in CM23 (Table 2). Among these 27 genes, five genes located on chromosome 9 have no homologue in *O. sativa* genome (Table 2). In agreement with the GSEA analysis (Fig. 7) showing alterations on carbon metabolism, photosynthesis and photorespiration, a homologous gene associated with alanine aminotransferase metabolism in rice, *OsALT-2*, OMERI07G00150, for a gene described in *O. sativa* as “Ribose-5-phosphate isomerase precursor” (OMERI07G03430) and for a glycolate oxidase (EC 1.1.3.15) (OMERI07G02060) were introgressed from *O. meridionalis* and down-regulated by +Fe in CM23 (Table 2).

## Discussion

Until now, no major locus conferring Fe excess tolerance has been fine-mapped or cloned (Meng *et al*., 2017). Selection of Fe tolerant rice genotypes is difficult for two reasons: (1) the quantitative nature of the trait controlled by multiple small and medium effect loci (Wu *et al*., 2014, Diop *et al*., 2020); and (2) the narrowed genetic diversity of the domesticated *O. sativa* varieties (Jacquemin *et al*., 2013, Menguer *et al*., 2017). To overcome these limitations, ILs and CSSLs using RWR as donor parents can be useful (Arbelaez *et al*., 2015, Bierschenk *et al*., 2020). The use of a population of interspecific IL derived from crosses between *O. sativa* and *O. meridionalis* (Arbelaez *et al*., 2015) was justified by previously reported tolerance of *O. meridionalis* accessions to Fe toxic conditions (Bierschenk *et al*., 2020). This species’ tolerance is probably associated with adaptation to its natural habitat in Northern Australia (Vaughan, 1994, Juliano *et al*., 2005), a region characterized by wetlands and Fe rich soils (Cogle *et al*., 2011). New QTLs were identified on chromosomes 1, 4, and 9 (Table 1). QTLs for LBS and relative decrease in shoot DW were also identified in different regions of chromosome 1 and 4 in previous studies (Wu *et al*., 1997, Wu *et al*., 1998, Dufey *et al*., 2009, Quinet *et al*., 2012, Wu *et al*., 2014). However, this is the first report showing that Fe tolerance traits from *O. meridionalis* can be introgressed into *O. sativa*. Our results highlight the importance of ILs as a tool for genetic studies in interspecific crosses of rice (Guo and Ye, 2014, Arbelaez *et al*., 2015).

Despite the fact that the *O. meridionalis* parent had low shoot Fe concentrations pointing to an exclusion mechanism, our physiological data suggested a shoot-based Fe tolerance mechanism in its progeny CM23. The interspecific hybrid exhibited high Fe storage in leaf sheaths and culms but less in the photosynthetically more active leaf blade (Fig. 4), a mechanism that was previously suggested to confer Fe tolerance (Audebert, 2006). Since translocation of Fe via the xylem follows the transpiration stream (Becker and Asch, 2005), the high Fe concentration in shoots of CM23 might be associated with the elevated transpiration rate of this genotype (Fig. S3D). CM23 also showed lower MDA levels compared to *O. sativa* (Fig. 3B-C), indicating that Fe is chelated or compartmentalized to avoid toxicity. Our previous work suggested shoot-based tolerance to +Fe in *O. sativa* can be achieved by modulating ascorbate recycling (Wu *et al*., 2017), but we did not find indications that the same mechanisms operates in CM23 (data not shown). Taken together, our results demonstrate the importance of introgressed segments from *O. meridionalis* in shoot tolerance to +Fe. suggesting that the *O. meridionalis* parent possesses novel shoot tolerance mechanisms in addition to its Fe exclusion capacity and pyramids these complementary tolerance mechanisms.

Several studies reported the regulation of genes in response to Fe deficiency in cultivated and wild rice (Ishimaru *et al*., 2009, Zheng *et al*., 2009, Bashir *et al*., 2014, Wairich *et al*., 2019), while others have explored mechanism and genes regulated by excess Fe (Quinet *et al*., 2012, Bashir *et al*., 2014, Stein *et al*., 2019). However, to the best of our knowledge, this is the first study to explore transcriptional responses of a wild rice hybrid and its parents to Fe toxicity. Since the genome of CM23 has not been sequenced, we reconstructed the interspecific hybrid genome *in silico* to map the sequences from the transcriptomic analysis. This approach illustrated that the CM23 transcriptome was less responsive to Fe compared to the parents, and functional categorization of DEGs showed striking differences (Fig. 6 and 7). Given the increasing importance of crop wild relatives as sources of novel traits for agricultural crops (Castaneda-Alvarez et al., 2016), we suggest that our methodological approach might be useful for other similar studies investigating transcriptomic regulation in crops and their wild relatives.

The plausibility of our transcriptome data is confirmed by the regulation of well characterized Fe-responsive genes. Two rice ferritin genes (*OsFER1* and *OsFER2*) were up-regulated in *O. sativa* and CM23 under +Fe. Ferritins are important proteins storing Fe within the cell, and their accumulation has been shown in rice (Silveira *et al*., 2009, Stein et al 2009), maize (Briat *et al*., 2010) and Arabidopsis (Tarantino *et al*., 2010) in response to Fe overload. In addition, VIT-like genes, Os04g0538400 and OsVIT2 (Os09g0396900), involved in the transport of Fe across the tonoplast (Zhang *et al*., 2012) were up-regulated under Fe stress. These genes were also differentially regulated in similar previous experiments in cultivated rice (Quinet *et al*., 2012, Aung *et al*., 2018, Stein *et al*., 2019). We also identified well-characterized genes involved in Fe-homeostasis that were down-regulated in our dataset. *OsIRO2*, one of the positive regulators of Fe deficiency response (Ogo *et al*., 2006), was down-regulated in all genotypes, suggesting all of them attempted to decrease Fe uptake. These data suggest that these genes can be considered as marker genes for Fe toxicity stress.

However, the main purpose of this study was not to identify Fe responsive genes but rather genes that may be involved in the shoot based tolerance mechanism of CM23, such as genes exclusively differentially regulated in CM23. Although the number of these genes was abundant, and it is possible that genes with hitherto unknown function conferred tolerance, it is worthwhile discussing genes with an annotation related to metal homeostasis. Among those were two metallothionein-like proteins. These proteins buffer cytosolic metal concentration by the formation of mercaptide bonds between the metal and their numerous Cys residues (Kumar *et al*., 2012).

Another category that could be of particular importance in conferring tolerance to CM23 are differentially regulated genes introgressed from *O. meridionalis* into CM23 (Table 8). Among those were GSTs, which are multifunctional enzymes playing a role in cellular detoxification (Salinas and Wong, 1999). In a previous microarray study, GSTs were also differentially regulated by +Fe in shoots of the tolerant *O. sativa* genotype FL483 compared to its more sensitive parent (Wu *et al*., 2017). Here we found that four Fe-responsive GSTs from *O. meridionalis* introgressed into CM23 (one in chromosome 7 and three closely located genes in chromosome 10) were up-regulated in response to +Fe. Also, GSTs were previously suggested as candidate genes conferring tolerance to +Fe in genotypes of *O. sativa* in a genome wide association study (Matthus *et al*., 2015). Within the chromosome 9 introgression segment, where our main QTL was located, only 15 genes were up-regulated by +Fe in CM23 (Table 2). Plausible candidate genes involved in +Fe response include also one ABC transporter (OMERI09G04710) and one MFS transporter (OMERI09G06320). ABC transporters were shown to detoxify As and Cd by transporting them into vacuoles (Zhang *et al*., 2018, Fu *et al*., 2019), whereas MFSs were involved in Fe homeostasis as transporters of nicotianamine and deoxymugineic acid (Che *et al*., 2019, Haydon *et al*., 2012). In order to confirm the involvement of these genes in Fe tolerance, further experiments would be required such as fine mapping and map-based cloning of QTL, or reverse genetic studies including genome editing. Overall our physiological and transcriptomic data suggest a shoot-based tolerance mechanism in the interspecific hybrid CM23, which is partly explained by retention of Fe in the leaf blade and culm, and could be regulated by genes involved in iron chelation, detoxification and partitioning.

In conclusion, our results highlight that RWR can be a rich reservoir of novel traits allowing plants to adapt to abiotic stress conditions. Introgressions from *O. meridionalis* conferred exceptional Fe tolerance to the interspecific hybrid line despite its very high shoot Fe concentrations. We further demonstrated that the availability of genome sequences for the wild relatives of crops form a powerful resource that allows for comparative transcriptional analyses of different species and crosses between them. This approach provided us with plausible candidate genes involved in Fe tolerance originating from a RWR that remain to be explored further.

## Supplementary data

Figure S1: Physiological parameters evaluated in the screening of 32 introgression lines under Fe excess (250 mg L^-1^ Fe^2+^) after fifteen days of treatment

Figure S2: Relative carotenoid reflectance index I and II in plants of *O. sativa* cv. Curinga, *O. meridionalis* (Ng. acc. W2112) and CM23 exposed to Fe toxicity (250 mg L^-1^ Fe^2+^)

Figure S3: Responses of gas exchange in plants of *O. sativa* cv. Curinga, *O. meridionalis* (Ng. acc. W2112) and CM23 exposed to Fe toxicity (250 mg L^-1^ Fe^2+^ for 8 and 13 days)

Table S1: Traits evaluated in screening of CM population under Fe toxicity

Table S2: Transcriptome data sets with genes differentially expressed in shoots of *Oryza sativa* cv. Curinga after seven days under Fe excess (250 mg L^-1^ Fe^2+^)

Table S3: Transcriptome data sets with genes differentially expressed in shoots of *Oryza meridionalis* (Ng. acc. W2112) after seven days under Fe excess (250 mg L^-1^ Fe^2+^)

Table S4: Transcriptome data sets with genes differentially expressed in shoots of IL CM23 after seven days under Fe excess (250 mg L^-1^ Fe^2+^)

Table S5: Transcriptome data sets with homologue inferred genes commonly up-regulated in shoots of *O. sativa, O. meridionalis* and IL CM23 after seven days under Fe excess (250 mg L^-1^ Fe^2+^)

Table S6: Transcriptome data sets with homologue inferred genes commonly down-regulated in shoots of *O. sativa, O. meridionalis* and IL CM23 after seven days under Fe excess (250 mg L^-1^ Fe^2+^)

Table S7: Genes differentially expressed regulated commonly between CM23 and O. sativa cv. Curinga and logFC > 2.0 and logFC < 2.0

Table S8: Genes exclusively differentially expressed in CM23 (when compared with O. sativa cv. Curinga)

## Abbreviations

+Fe: Fe toxicity
AAS: Atomic Absorption Spectrometry
CRI: carotenoid reflectance index
CSSL: chromosome segment substitution lines
DEG: differentially expressed gene
DW: dry weight
IL: introgression line
LB: leaf blades
LBS: leaf bronzing score
MDA: malondialdehyde
NDVI: normalized difference vegetation index
QTL: quantitative trait *loci*
RDVI: renormalized difference vegetation index
ROS: reactive oxygen species
RWR: rice wild relatives
TF: transcription factor

## Acknowledgements

This study was financed in part by the Coordenação de Aperfeiçoamento de Pessoal de Nível Superior-Brasil (CAPES)-Finance Code 001 and Conselho Nacional de Desenvolvimento Científico e Tecnológico (CNPq), which granted fellowships to AW, BHNO, JPF, MMP and FKR. We also thank FAPERGS (Pronex process 16/25.51-0000493-5) and CAPES/DAAD/Probral (process 88881.144076/2017-01). We also thank Prof. Susan McCouch for providing seeds of the interspecific introgression lines.

